# Urinary multi-omics reveal non-invasive diagnostic biomarkers in clear cell renal cell carcinoma

**DOI:** 10.1101/2024.08.12.607453

**Authors:** Gustav Jonsson, Maura Hofmann, Tiago Oliveira, Ursula Lemberger, Karel Stejskal, Gabriela Krššáková, Irma Sakic, Maria Novatchkova, Stefan Mereiter, Gerlinde Grabmann, Thomas Köcher, Zeljko Kikic, Gerald N. Rechberger, Thomas Züllig, Bernhard Englinger, Manuela Schmidinger, Josef M. Penninger

## Abstract

Clear cell renal cell carcinoma (ccRCC) is the kidney malignancy with the highest incidence and mortality rates. Despite the high patient burden, there are no biomarkers for rapid diagnosis and public health surveillance. Urine would be an ideal source of ccRCC biomarkers due to the low invasiveness, easy accessibility, and the kidney’s intrinsic role in filtering urine. In the present work, by combining proteomics, lipidomics and metabolomics, we detected urogenital metabolic dysregulation in ccRCC patients with increased lipid metabolism, altered mitochondrial respiration signatures and increased urinary lipid content. Importantly, we identify three early-stage diagnostic biomarkers for ccRCC in urine samples: Serum amyloid A1 (SAA1), Haptoglobin (HP) and Lipocalin 15 (LCN15). We further implemented a parallel reaction monitoring mass spectrometry protocol for rapid and sensitive detection of SAA1, HP and LCN15 and combined all three proteins into a diagnostic UrineScore. In our discovery cohort, this score had a performance accuracy of 96% in receiver operating characteristic curve (ROC) analysis for classification of ccRCC versus control cases. Our data identifies tractable and highly efficacious urinary biomarkers for ccRCC diagnosis and serve as a first step towards the development of more rapid and accessible urinary diagnostic platforms.

## Introduction

Renal cell carcinomas (RCCs) are a heterogenous group of cancers arising from the proximal convoluted tubule of the kidney. Many histological subtypes of renal cell carcinomas have been described, with the three most common ones being clear cell renal cell carcinoma (ccRCC), papillary renal cell carcinoma (pRCC) and chromophobe renal cell carcinoma (chRCC). Out of these three subtypes, ccRCC is the most common, making up approximately 80% of all RCC cases, and also the deadliest [1, 2]. ccRCC is primarily driven by loss-of-function mutations or epigenetic silencing of the *Von Hippel Lindau (VHL)* tumor suppressor, leading to the initiation of a genetic hypoxia program which drives tumor progression [3, 4]. Despite this clearly identified mechanism, solely screening for *VHL* mutations for disease detection is not sufficient since not all patients present with *VHL* gene mutations. Furthermore, multiple additional co-drivers have also been identified, such as PBRM1, SETD2, KDM5C, or BAP1 [5, 6], convoluting genetic-based disease screening.

Despite significant advances in the treatment of ccRCC using small molecule inhibitors and immunotherapy [7–9], early screenable markers for diagnosis are still lacking and most patients are diagnosed through computed tomography when a tumor is already suspected, for example due to palpable renal masses or hematuria [10]. Hematuria is a very common sign of ccRCC, and other renal cell malignancies, but in and of itself insufficient for a reliable diagnosis. The proximal convoluted tubule, where ccRCC arises, is responsible for secreting and reabsorbing solutes between blood and urine [11] increasing the likelihood that secreted or shed biomarkers are present and detectable in the urine. Furthermore, ccRCC predominantly affects people of the age of 60 and upwards, and very rarely younger individuals [10]. In many countries, the older population is already part of population-based screening programs and health check-ups in which urine is routinely donated and analyzed, making urine samples an ideal source for early detection of ccRCC. Urine also allows easy longitudinal sampling and is minimally invase. Omics approaches have been used on urine samples from renal cell carcinomas in the past for the purpose of biomarker discovery, predominantly metabolomics due to the natural abundance of high levels of excreted metabolites as waste products in urine [12]. However, multiple omics approaches have yet to be integrated on the same urine samples to draw multi-modal conclusions about the dysregulation of the urinary landscape in ccRCC, and how this can be exploited for biomarker discovery.

In the present work, by combining proteomics, lipidomics and metabolomics, we detected urogenital metabolic dysregulation in ccRCC patients with increased lipid metabolism, mitochondrial respiration signatures and increased urinary lipid content. We further aimed to explore urinary biomarkers to reliably detect ccRCC. In a clinical cohort of controls, ccRCC patients and non-clear renal cell carcinoma (nccRCC) patients, we discovered and validated three proteins as ccRCC diagnostic biomarkers: SAA1, HP, and LCN15. We further developed a parallel reaction monitoring mass spectrometry (PRM-MS) signature for rapid and sensitive quantitative detection of these three marker proteins in patient urine, discriminating between controls and ccRCC patients with a performance accuracy of 96%.

## Results

### Urine from ccRCC patients is indicative of metabolic dysregulation

To characterize the landscape of ccRCC urine and to potentially derive biomarkers for the disease we initially performed proteomics on urine sediment and supernatant on a cohort of 22 ccRCC patients and 12 controls **(Fig. 1A, Supplementary Table 1**). Of note, in pilot studies we found that precipitating proteins in two steps from the urine samples (tested on supernatant) starting with an acetone precipitation followed by a chloroform:methanol precipitation yielded the highest number of unique peptides and proteins identified (**Supplementary Fig. 1**). This method was subsequently used for all protein preparations. All discovery-phase proteomics analysis was done through data independent acquisition (DIA). Gene ontology (GO) term analysis of all upregulated proteins in ccRCC urine supernatants from proteomics, compared to controls, predominantly revealed metabolic dysregulation affecting lipid metabolism and function through, e.g. fatty acid transport and high-density lipoparticle remodeling (**Fig. 1B**, **Supplementary Table 2**). Further, GO term analysis on sediment proteomics from control and ccRCC patient urine samples showed that mitochondrial respiration and electron transport processes were upregulated in ccRCC patients (**Fig. 1C**, **Supplementary Table 3**).

**Figure 1.**
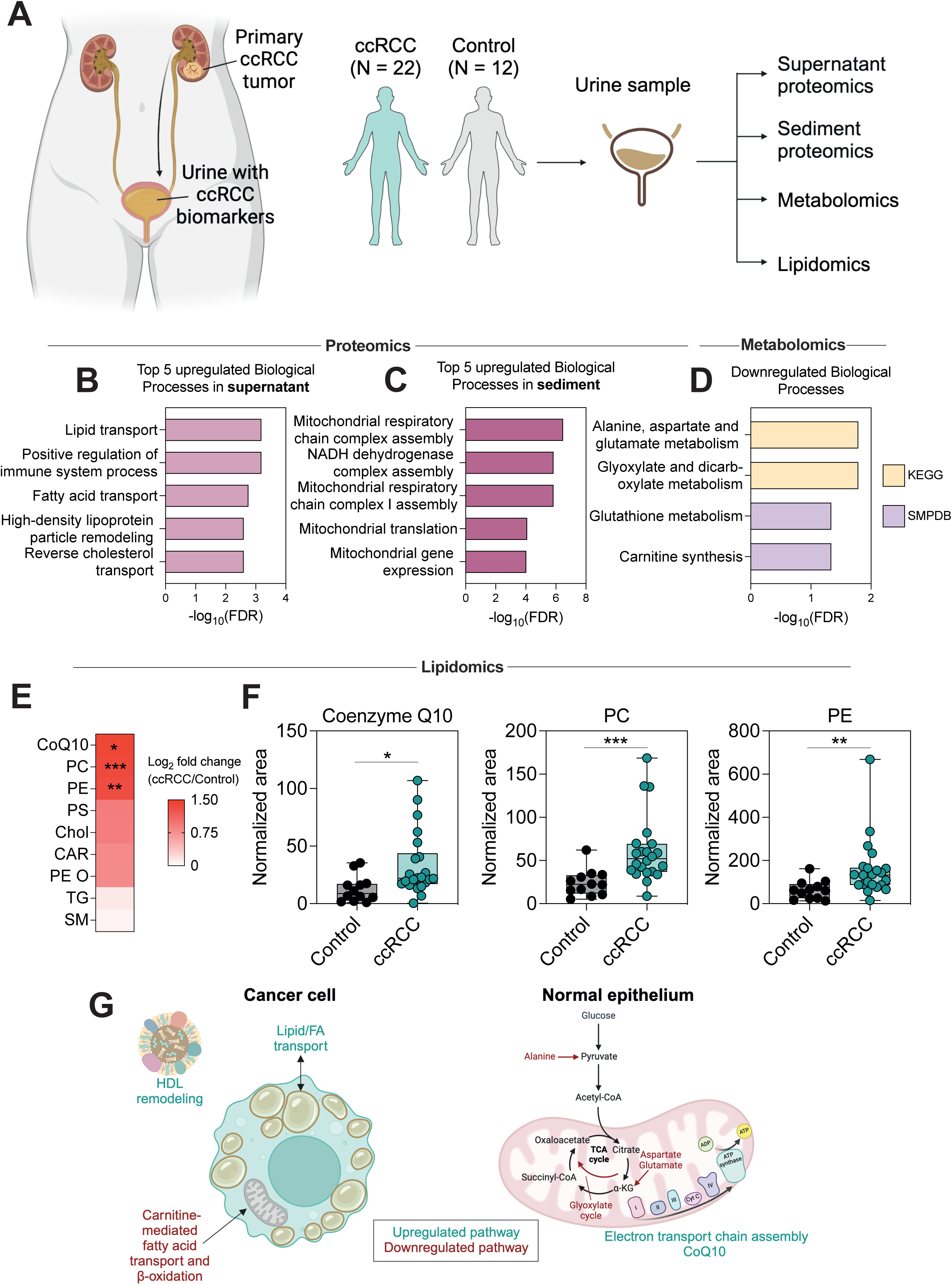
Multi-omics analysis of ccRCC patient urine indicates metabolic dysregulation. **A.** Schematic of detection of urine analytes in ccRCC patients passing through the urogenital tract and accumulating in the urine bladder before discharge in a ccRCC patient cohort. Schematic created with BioRender.com. **B.** Gene ontology (GO) term analysis using Enrichr on all upregulated proteins in ccRCC urine supernatant compared to Control urine supernatant. FDR = False discovery rate. **C.** Gene ontology (GO) term analysis using Enrichr on all upregulated proteins in ccRCC urine sediment compared to Control urine sediment. FDR = False discovery rate. **D.** Gene ontology (GO) term analysis using Metaboanalyst on all downregulated metabolites in ccRCC urine supernatant compared to Control urine supernatant. The downregulated metabolites were matched against two different databases: KEGG and SMPDB. FDR = False discovery rate. **E.** Heatmap showing fold change between the detected lipid families in ccRCC patient urine and control urine. CoQ10 = Coenzyme Q10, PC = Phosphatidylcholines, PE = Phosphatidylethanolamine, PS = Phosphatidylserine, Chol = Cholesterol, CAR = Acylcarnitines, PE O = Ether-linked phosphatidylethanolamine, TG = Triglyceride, SM = Sphingomyelin. **F.** Boxplot of individual values for the three significantly higher lipid families from **E**. CoQ10 = Coenzyme Q10, PC = Phosphatidylcholines, PE = Phosphatidylethanolamine. * = p < 0.05, ** = p < 0.01, *** = p < 0.001. **G.** Schematic of proposed altered metabolic network in the urogenital tract of ccRCC patients. Green represents upregulated pathways and red represents downregulated pathways. HDL = High-density lipoprotein, CoA = Coenzyme A, α-KG = alpha-ketoglutarate, CoQ10 = Coenzyme Q10. Schematic created with BioRender.com.

Since the proteomics data indicated dysregulated metabolism throughout the urogenital tract, we further performed metabolomics and lipidomics on the urine samples. Untargeted metabolomics predominantly revealed downregulated metabolites in ccRCC patients (**Supplementary Fig. 2A**). After filtering the hits further to only include metabolites from the mzCloud database and our in-house validated database, 36 downregulated metabolites remained (**Supplementary Fig 2A-B, Supplementary Table 4**); some examples of which are shown in **Supplementary Fig. 2C**. GO term analysis on the 36 downregulated metabolites, matched against the KEGG and SMPDB databases, revealed downregulation of (i) aspartate, alanine and glutamate metabolism, (ii) glyoxylate metabolism, (iii) glutathione metabolism and (iv) carnitine synthesis (**Fig. 1D**).

Our lipidomics analysis revealed that 3 out of 9 detected lipid classes were upregulated in ccRCC compared to controls: Coenzyme Q10, phosphatidylcholines (PCs) and phosphatidylethanolamines (PEs) (**Fig. 1E-F**, **Supplementary Table 5**). Combined, most of the measured individual PC and PE species were significantly upregulated in ccRCC compared to controls (**Supplementary Fig. 3A-B**). In summary, the proteomics, metabolomics and lipidomics data indicate dysregulated metabolism with increased electron transport chain activity and increased lipid metabolism throughout the urogenital tract, summarized in **Fig. 1G**. The observed lipid phenotype is likely linked to cancer cells whereas electron transport chain phenotypes are linked to healthy, shed epithelium in the urine sediment. ccRCC cells are known to be lipid laden and accumulate high levels of lipids. Increased PC and PE levels in the urine likely come from increased lipid transport in these cells. Further, carnitine is important for transport of fatty acids through the mitochondrial membrane, leading to their breakdown in β-oxidation, meaning that the reduction of carnitine is likely to contribute to the increased lipid content in the cells. Combined, the lipid landscape of the urine supernatant appears to be representative of what is going on in the tumors. ccRCC is known to be independent of oxidative phosphorylation [13] indicating that upregulated respiratory chain pathways in the urine sediment most likely come from increased shedding of healthy epithelium or other shed urogenital cells, and not actually from the tumor itself. We further detected upregulation of CoQ10 which is important for electron chain transport function, and downregulation of alternative entry points into the TCA cycle (alanine, aspartate and glutamate metabolism, and glyoxylate metabolism) (**Fig. 1E-G**), indicating that conventional glycolysis is used for increased oxidative phosphorylation.

### Lipidomics and metabolomics provide putative diagnostic biomarkers for ccRCC

ccRCC is the deadliest type of renal carcinoma (**Fig. 2A**) with a tendency to appear asymptomatic at early stages leading to complications at the later stages of disease. Compared to many other malignancies, ccRCC lacks liquid biopsy biomarkers for early diagnosis and prognosis [14, 15]. We hypothesized that our multi-omics dataset could lead to the identification of reliable early-stage biomarkers since a majority of our ccRCC cohort, included in the analyses, consisted of early pT1 stage tumors (**Fig. 2B**). Initially we assessed the ability of the three significantly upregulated lipid classes (CoQ10, PC and PE) to distinguish between ccRCC and controls in an area under the receiver operating characteristic curve (AUROC) analysis (**Fig. 2C**). CoQ10, PC and PE all had a good performance of AUROC = 0.7992, AUROC = 0.8826 and AUROC = 0.8258, respectively, in distinguishing between ccRCC and control cases. Furthermore, when we assessed all significantly upregulated individual lipid species, all of them had an AUROC > 0.7, with PC 38:3 and PC 38:6 having the highest AUROC of 0.9129 and 0.8996, respectively (**Fig. 2D**), indicating the potential use of PC family members in ccRCC diagnosis.

**Figure 2.**
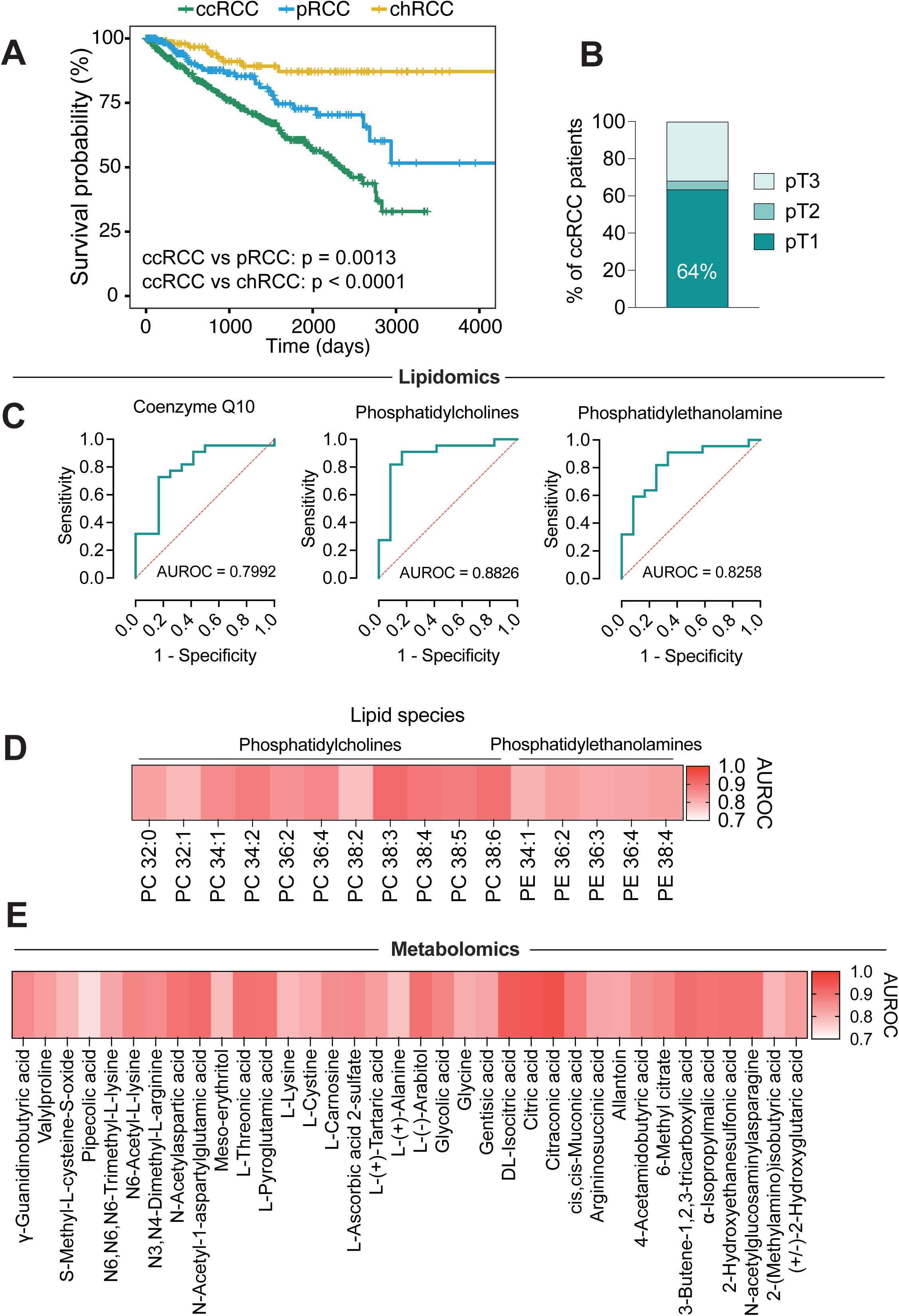
Lipidomics and metabolomics reveal biomarkers that distinguish between controls and ccRCC patients. **A.** Survival curves ccRCC, pRCC and chRCC patients present in the TCGA database. TCGA = The Cancer Genome Atlas, ccRCC = clear cell renal cell carcinomas, pRCC = papillary renal cell carcinomas, chRCC = chromophobe renal cell carcinoma. Statistical significance assessed through Log-rank tests. **B.** Distribution of tumor stages at the timepoint of presentation in the clinic for ccRCC patients. Abbreviations: pT1 = Tumor stage 1. The tumor is a maximum of 7 cm across. pT2 = Tumor stage 2. The tumor is larger than 7 cm across. pT3 = Tumor stage 3. The tumor has grown into a major renal vein (e.g. vena cava or renal vein) or into neighboring tissues but has not spread past Gerota’s fascia or into the adrenal gland. **C.** Assessment of the ability of the three significantly upregulated lipid families (CoQ10, PC and PE, respectively) in distinguishing between ccRCC and control cases using AUROC analysis. CoQ10 = Coenzyme Q10, PC = Phosphatidylcholines, PE = Phosphatidylethanolamine, AUROC = Area under receiver operating characteristic curve. **D.** Heatmap showing AUROC values for individual lipid species from PC and PE that were upregulated in ccRCC compared to controls. PC = Phosphatidylcholines, PE = Phosphatidylethanolamine, AUROC = Area under receiver operating characteristic curve. **E.** Heatmap showing AUROC values for individual metabolites that were downregulated in ccRCC compared to controls. AUROC = Area under receiver operating characteristic curve.

Similar trends were observed when assessing downregulated metabolites. All significantly downregulated metabolites presented with an AUROC > 0.7 in being able to distinguish between ccRCC and controls, with citraconic acid, citric acid and DL-Isocitric acid scoring the highest AUROC values of 0.9545, 0.9432, and 0.9356, respectively (**Fig. 2E**). These data indicate that altered lipid and metabolite signatures could be used to distinguish ccRCC from healthy controls in urine.

### Proteomics provides high-confidence diagnostic biomarkers for ccRCC

Despite the statistical validity of lipids and metabolites in distinguishing between ccRCC and control cases and generating interesting biological data, for biomarker utilization they do come with a set of challenges. For lipidomics, it is difficult to achieve complete analytical coverage and accurate abundancy quantification remains an issue [16]. Furthermore, in our study the total number of identified lipid species was on the lower side and all species were not identified in every single sample. For metabolomics, we only detected significantly downregulated metabolites in ccRCC patients. Urine as a sample has low levels of analytes, and basing diagnosis on downregulation of already lowly abundant analytes is challenging.

For these reasons, we focused our efforts at finding an easily assessable biomarker on proteomics. Mass spectrometry-based proteomics are frequently available and established in hospital diagnostic laboratories compared to lipidomics and metabolomics. Protein level diagnosis further offers the benefit of being able to develop ELISA based approaches for cost-effective and routine disease screening. Our bulk proteomics data on both urine sediment and supernatant revealed upregulated proteins with potential as diagnostic biomarkers; e.g. NAT10, APOL1, NDUFS7 and MRPL48 for sediment (**Fig. 3A**) and SAA1, HP and LCN15 for supernatant (**Fig. 3B**). Mass spectrometry data is available in **Supplementary Table 2-3**. For any potential diagnostic biomarker, in addition to being able to distinguish between ccRCC and controls, we also wanted to test if such markers can distinguish between ccRCC and non-clear cell renal cell carcinomas (nccRCC), such as pRCC and chRCC. For this purpose, we extended our cohort to also include 9 nccRCC patients, from which 8 patients had pRCC and 1 patient chRCC (**Fig. 3C**, **Supplementary Table 1**). ccRCC patients had the highest amounts of protein in the urine supernatant (**Supplementary Fig. 4A**). The overall amount of protein found in the urine of ccRCC patients was independent of the tumor stage (**Supplementary Fig. 4B**). Furthermore, for leukocytes and inflammatory damage markers (Creatinine and C-reactive protein) in the serum, no differences were found between ccRCC and nccRCC patients, nor between different stages of ccRCC (**Supplementary Fig. 4C, D**).

**Figure 3.**
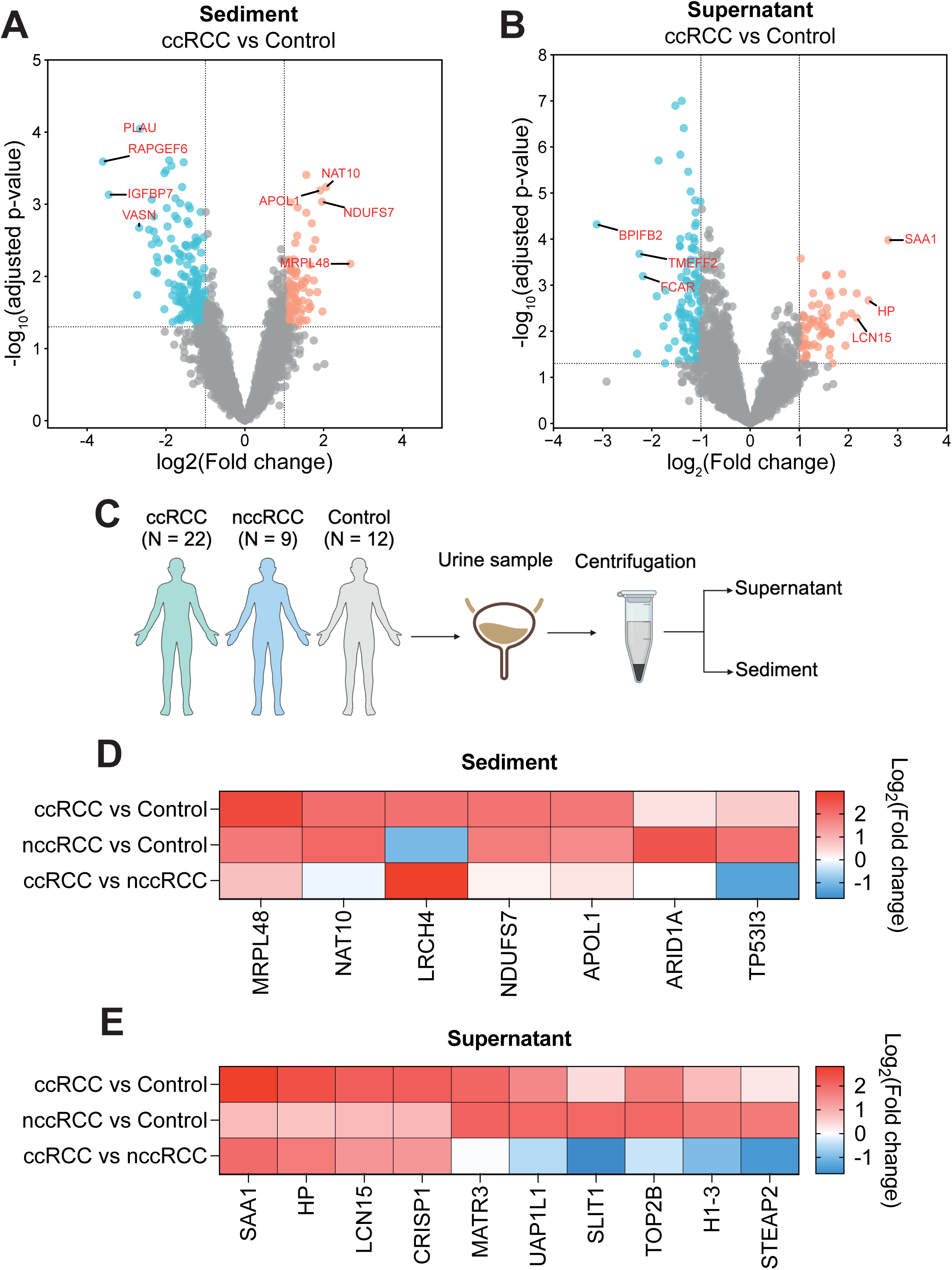
Identification of protein biomarkers for ccRCC diagnosis. **A.** Volcano plot showing upregulated (red) and downregulated (blue) proteins in ccRCC patients compared to controls in urine sediment. Adjusted p values calculated via limma-moderated Benjamini–Hochberg-corrected two-sided t-test. FC = fold change. **B.** Volcano plot showing upregulated (red) and downregulated (blue) proteins in ccRCC patients compared to controls in urine supernatant. Adjusted p values calculated via limma-moderated Benjamini–Hochberg-corrected two-sided t-test. FC = fold change. **C.** Schematic of sample acquisition from ccRCC patients, nccRCC patients and controls for urine supernatant and sediment proteomics. **D.** Heatmap of the highest upregulated proteins in ccRCC vs control urine (top row) and nccRCC vs control urine (middle row) in urine sediments. A comparison between ccRCC and nccRCC is also shown in the bottom row. **E.** Heatmap of the highest upregulated proteins in ccRCC vs control urine (top row) and nccRCC vs control urine (middle row) in urine supernatans. A comparison between ccRCC and nccRCC is also shown in the bottom row.

When comparing nccRCC urine sediments to controls, NAT10 and MRPL48 were the highest upregulated proteins, just as for ccRCC urine sediments compared to controls (**Fig. 3D**, **Supplementary Fig. 5A**). Overall, the upregulated protein profiles were similar for both ccRCC and nccRCC, with LRCH4 and TP53I3 being the only proteins with the potential of distinguishing between ccRCC and nccRCC (**Fig 3D**, **Supplementary Fig. 5B**). For supernatants however, different proteins were upregulated between cancer and control for ccRCC and nccRCC, except for MATR3 (**Fig. 3E**, **Supplementary Fig. 5C-D**). Due to the different urinary protein profiles of ccRCC and nccRCC, urine supernatant proteins were chosen as the preferred source of diagnostic biomarkers. Furthermore, reasonable protein quantities for proteomics could not be extracted from all samples (**Supplementary Table 3**). Serum Amyloid A1 (SAA1), Haptoglobin (HP) and Lipocalin 15 (LCN15) were selected as the putative biomarkers based on the following criteria: (i) having a log_2_(Fold change) > 2 in ccRCC compared to controls, (ii) upregulated in the ccRCC and control comparison, and not the nccRCC and control comparison which eliminates MATR3 and (iii) not being male or female specific, e.g. CRISP1 is male specific.

### Parallel reaction monitoring mass spectrometry for rapid detection of SAA1, HP and LCN15 in ccRCC urine

Conventional bulk proteomics allows discovery at the full proteome level but is not suitable for rapid diagnostics and population screenings in a clinical setting due to long instrument run times, costs, and computationally intensive data analysis. Therefore, we utilized parallel reaction monitoring mass spectrometry (PRM-MS) to develop a tractable diagnostic modality for SAA1, HP and LCN15. PRM-MS is an ion monitoring technique allowing parallel high-resolution detection of peptides of interest, drastically reducing the run time per sample and increasing specificity compared to bulk proteomics and other ion monitoring techniques [17].

In our PRM-MS approach, we included all peptides for SAA1, HP and LCN15 detected by our bulk proteomics approach with clearly defined elution patterns (**Supplementary Table 6**). For standardization, we included peptides from three normalization proteins, namely Uromodulin (UMOD), Kallikrein-1 (KLK1) and Apolipoprotein D (APOD). A normalization protein was defined as a protein which was found in every sample in all three groups (Control, nccRCC and ccRCC) and with similar expression levels across all three experiment groups per protein (**Supplementary Table 2**). The peptide area was calculated for every selected peptide and the PRM score for each diagnostic protein was determined by dividing the sum of all peptide areas from individual proteins of interest (SAA1, HP, LCN15) with the sum of all peptides from all three normalization proteins (**Fig. 4A**).

**Figure 4.**
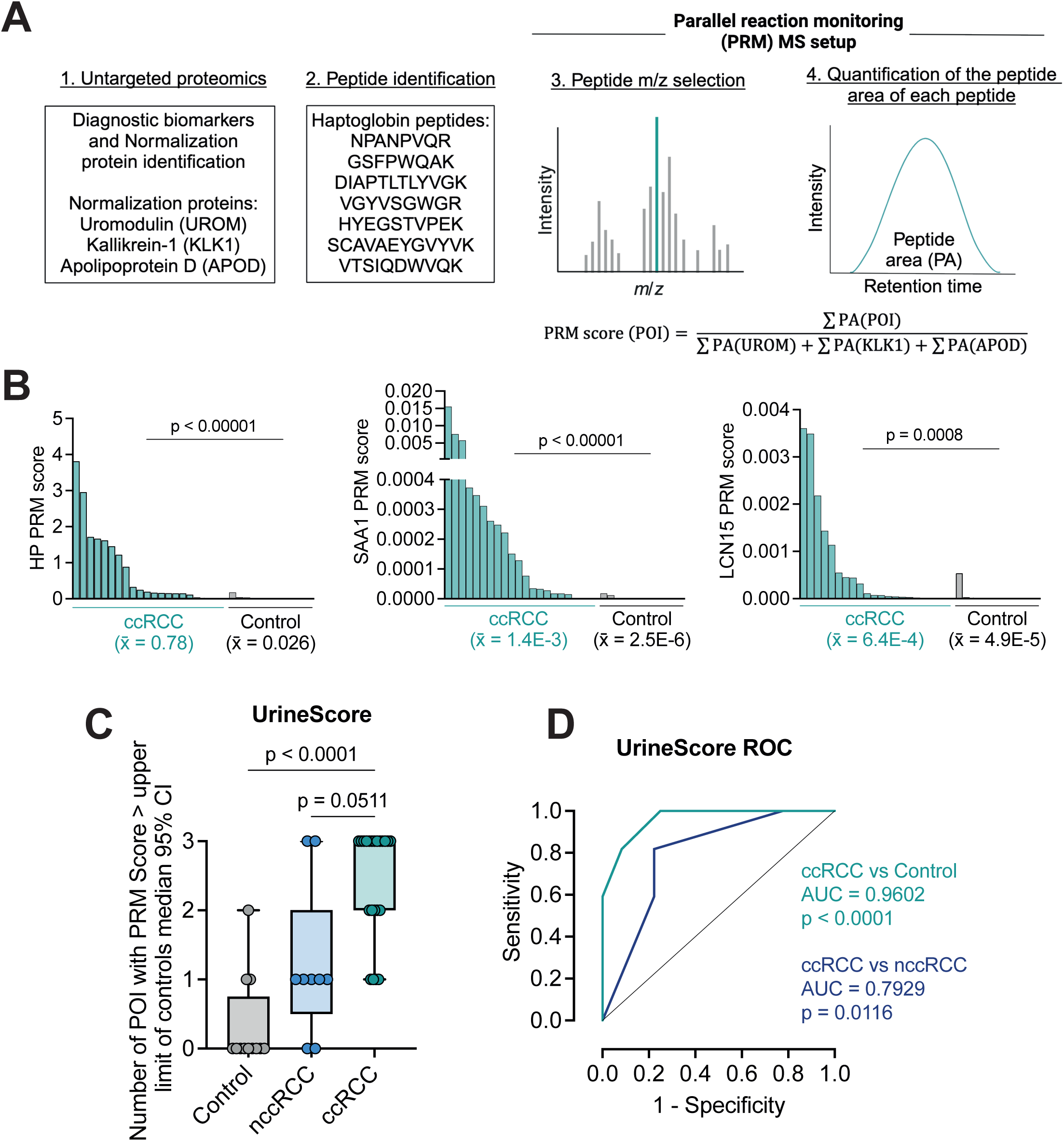
Parallel reaction monitoring mass spectrometry accurately diagnoses ccRCC. **A.** Schematic of the PRM-MS method used to quantify the SAA1, HP and LCN15 levels in urine samples. Levels are determined based on normalization to three normalization proteins (UMOD, KLK1 and APOD). The peptide area for each detected peptide from a specified protein is quantified, detected peptides from Haptoglobin are shown as an example. Abbreviations: PA = Peptide area, POI = Protein of interest (SAA1, HP or LCN15). Schematic created using BioRender.com. **B.** Waterfall plot of PRM score for Haptoglobin, SAA1 and LCN15, respectively, compared between ccRCC and Control cohorts. Statistical significance assessed through Mann-Whitney U-test. x̄ = sample mean. **C.** Box plot of UrineScore for controls, nccRCC and ccRCC patients. Per protein of interest (POI) per patient or control, a value of 1 is derived if the PRM score is higher than the upper limit of the 95% CI for the control. Statistical significance assessed through Kruskal-Wallis test with Dunn’s multiple comparisons test. **D.** Receiver operating characteristic curve to assess the performance of the UrineScore in differentiating between ccRCC and control samples, and between ccRCC and nccRCC samples. Performance quantified through Area under the receiver operating characteristic curve (AUROC) analysis.

The PRM scores for SAA1, HP and LCN15 were all significantly higher in ccRCC patients compared to controls (**Fig. 4B**). The classification performance of each PRM score was calculated by the AUROC method and was evaluated to be 0.88, 0.89 and 0.84 for SAA1, HP and LCN15, respectively (**Supplementary Fig. 6**).

Lastly, all three individual PRM scores were combined into a cumulative UrineScore, detailed in the Methods section. In brief, the 95% confidence interval (CI) of the median was calculated for all PRM parameters for the control samples. Subsequently, each sample from the control, nccRCC and ccRCC groups was attributed a score of 1 per protein (SAA1, HP and LCN15) which is a value higher than the upper limit of the control 95% CI, meaning that the UrineScore is an integer between 0-3 per control or patient. This UrineScore was the highest for the ccRCC group (**Fig. 4C**) and had a very high-performance accuracy in distinguishing between control and ccRCC samples (AUROCC = 0.96), and also performed well between nccRCC and ccRCC samples (AUROCC = 0.79) (**Fig. 4D**). These data indicate that PRM-MS allows more rapid and sensitive diagnostic power compared to bulk proteomics, and that SAA1, HP and LCN15 can be used to diagnose ccRCC in our discovery cohort.

## Discussion

In 2020, approximately 431000 new cases of renal cell carcinomas were diagnosed worldwide [18]. Out of all of these cases, it is estimated that 80% of the renal cell carcinomas are ccRCC. With a growing incidence rate, there is increased interest in cost-effective and sensitive population-wide screening programs. Despite the high clinical interest, there are no validated biomarkers for clinical screening, and common diagnostic procedures such as computed tomography are not suitable for population-based screenings programs [19, 20].

Utilizing urinary proteomics, metabolomics and lipidomics, we uncovered aberrant metabolism throughout the urogenital tract of ccRCC patients. Our data suggests increased mitochondrial respiration and lipid metabolism. ccRCC is known to have a lipid rich cytoplasm [21]. We discovered increased lipid transport and a reduction in carnitine synthesis. Although our data comes from the urine and not directly from the tumor, one potential explanation for the increased lipid accumulation in ccRCC tumors could be a reduction of carnitine, which is essential for the breakdown of free fatty acids and potentially lipid transport to the tumor microenvironment, and not only increased lipid transport through the reverse cholesterol pathway in which cholesterol is recycled to the liver [22]. Amongst the individual lipid species detected, we noticed an increase in certain PE and PC species indicative of increased tissue damage and membrane shedding into the excreted compartment. We further found increased levels of CoQ10, corroborating our observations of increased mitochondrial respiration. Generally, ccRCC tend to be independent of oxidative phosphorylation as a source of energy [13], indicating that the mitochondrial respiration phenotypes we encountered in urine sediment are likely coming from shed epithelium and surrounding tissue damage from the tumor.

Proteomics on the urine supernatants proved the most reliable and feasible for diagnostic biomarker discovery. Bulk proteomics uncovered three putative diagnostic biomarkers (SAA1, HP and LCN15) in a small clinical cohort. Furthermore, through the establishment of PRM-MS for rapid and sensitive detection of these three proteins of interest we could derive PRM scores for SAA1, HP and LCN15 with high diagnostic power. When these PRM scores were combined into an overarching UrineScore, we calculated an overall performance in an AUROC analysis of 96% in distinguishing between healthy controls and ccRCC. The UrineScore also performed well in distinguishing between ccRCC and nccRCC patients, though this needs to be confirmed in larger patient cohorts.

The cellular source of SAA1, HP and LCN15 and how these three proteins are being filtered/shed into the urine remains to be answered. All three proteins are secreted proteins, meaning that they could directly be produced at high levels in the tumors and subsequently secreted into the urine. However, SAA1 and HP are also produced as acute phase proteins in the liver, and LCN15 is produced in the gastrointestinal tract. Future studies will have to show how large of a proportion of the markers found in the urine are directly derived from the tumor and/or other other cellular sources. It is possible that circulating serum proteins are found at higher levels in the urine of ccRCC patients due to tumor-induced tissue damage of the filtration unit of the kidney, effectively increasing the leakiness of serum proteins into the urine which is found in ccRCC associated hematuria. However, this does not appear to be a general mechanism since we in that case would likely have found a larger range of serum proteins as diagnostic markers.

The urine proteome of ccRCC patients has been studied before. For example, differences in urine protein content has been found amongst ccRCC patients depending on disease prognosis [23, 24] and venous infiltration [25]. Furthermore, attempts have been made at identifying diagnostic and prognostic protein biomarkers in the urine of ccRCC patients [26, 27]. However, none of these studies identified SAA1, HP or LCN15 in their analysis. Lastly, our three proteins of interest also differentiate between ccRCC and nccRCC, highlighting the specificity of these markers towards ccRCC. Validation trials need to be conducted to validate this readily screenable UrineScore in larger, independent clinical cohorts but the UrineScore highlights the potential of urine based diagnostic scoring not only for identifying renal cell carcinomas, but also for distinguishing between different histological subtypes.

## Materials and Methods

### Patient cohort

The study was approved by the ethical commission at the Medical University of Vienna (Ethik-Kommission Medizinische Universität Wien), study number 2224/2021, project title: Urinproteomik zur Validierung von Biomarkern für das klarzellige Nierenkarzinom – Pilotstudie (UrineProt).

Urine was collected from 40 patients who presented with a suspected primary renal mass at the Department of Urology of the Medical University of Vienna. Out of the 40 patients, 9 were excluded due to the renal mass being identified as something other than a renal cell carcinoma, such as oncocytomas, cysts, angiomyolipoma, papillary adenoma or a kidney-lodged metastasis. From the remaining 31 patients, 22 patients were characterized as ccRCC, 8 as pRCC and 1 as chromophobe RCC through histological assessment by a trained pathologist. For further analysis, the pRCC and chromophobe RCC patients were combined into one group of non-clear cell renal cell carcinoma (nccRCC) patients. Leukocytes, Creatinine and CRP levels were obtained in the routine blood analysis using standard methods at the Institute of Laboratory Medicine, Vienna General Hospital.

### Protein precipitation

3 ml of ccRCC patient, nccRCC patient or control urine were centrifuged at 1000xg for 10 min. The urine supernatant was isolated and mixed with 12 ml of acetone and incubated for a minimum of 2 hours at -20°C to allow for protein precipitation. Samples were centrifuged at 3000xg for 60 min and the supernatant discarded. The protein pellet was then resuspended in 100 μl of 8M Urea and a second precipitation step was performed to increase protein yields and to discard residual contaminants using chloroform and methanol [28]. 400 μl methanol was added to the protein solution and vortexed extensively. Subsequently, 200 μl of chloroform and 300 μl of water was added with vortexing steps in between. The mixtures were centrifuged at 10000xg for 15 min at room temperature to ensure phase separation. Following centrifugation, the top aqueous layer was removed leaving the lower chloroform fraction and white protein interface. To the remaining mixture, 400 μl of methanol was added followed by vortexing and centrifugation at 10000xg for 5 min. Supernatant was removed and the methanol wash was repeated twice. After the last methanol wash, the pellet was dried in a speed vacuum and resuspended in 8M Urea.

### Enzymatic digestion of precipitated proteins

Precipitated proteins were reduced in 10 mM DTT (Roche) at 37°C for 1 hour and diluted to 4M Urea. Following reduction, proteins were subsequently alkylated with 20 mM Indole-3-acetic acid (Merck) for 1h at room temperature in the dark and diluted to 2M Urea. Following reduction and alkylation, proteins were initially enzymatically digested with LysC (Lysyl Endopeptidase®, Mass Spectrometry Grade, FUJIFILM) at 37°C for 2hrs according to the manufacturer’s instructions. Lastly, peptides were further digested with trypsin (Trypsin Gold, Mass Spectrometry Grade, Promega) overnight at 37°C according to the manufacturer’s instructions.

### Bulk proteomics

#### Data-independent acquisition mass spectrometry

Individual peptide samples were analyzed by LC–MS/MS. The nano HPLC system used was an UltiMate 3000 nano HPLC RSLC (Thermo Scientific) equipped nano-electrospray source (CaptiveSpray source, Bruker Daltonics), coupled to a timsTOF HT mass spectrometer (Bruker Daltonics). Peptides samples were injected on a pre-column (PepMap C18, 5 mm × 300 μm × 5 μm, 100 Å pore size, Thermo Scientific) with 2% ACN/water (v/v) containing 0.1% TFA at a flow rate of 10 μL/min for 10 min. Peptides were then separated on a 25 cm Aurora ULTIMATE series HPLC column equipped with an emitter (CSI, 25 cm × 75 µm ID, 1.7 µm C18, IonOpticks) operating at 50°C and controlled by the Column Oven PRSO-V1-BR (Sonation), using UltiMate 3000 (Thermo Scientific Dionex). The analytical column flow was run at 300 nL/min with two mobile phases: water with 0.1% FA (A) and water with 80% acetonitrile and 0.08% formic acid (B). A and B were applied in linear gradients as follows (only B percentages reported): starting from 2% B: 2%-10% B in 10 min, 10%-24% B in 35 min, 24%-35% B in 15 min, 35%-95% B in 1 min, 95% for 5 min, and finally the column was equilibrated in 2% B for next 10 min (all % values are v/v; Water and ACN solvents were purchased from Thermo Fisher Scientific at LC-MS grade).

The LC system was coupled to a TIMS quadrupole time-of-flight mass spectromecter (timsTOF HT, Bruker Daltonics) and samples were measured in dia-PASEF mode. The CaptiveSpray source parameters were: 1600 V capillary voltage, 3.0 l/min dry gas, and 180 °C dry temperature. MS data was acquired in the MS scan mode, using positive polarity, 100-1700 m/z range, mobility range was set up from 0.64-1.42 V s/cm^2^, ramp time was set to 166 ms and the estimated cycle time was 1.52s. Collision energy was 20 eV at 1/K_0_ 0.6 V s/cm^2^, and 80 eV at 1/K_0_ 1.6 V s/cm^2^. Automatic calibration of ion mobility was enabled. The timsTOF HT was operated in DIA mode where 1 MS1 scan was followed by 8 DIA-PASEF frames.

#### Proteomics data analysis

DIA data was analyzed in Spectronaut 18.5 [29] (Biognosys). Trypsin/P was specified as a proteolytic enzyme and up to 2 missed cleavages were allowed in the Pulsar direct DIA search. Dynamic mass tolerance was applied for the TOF calibration. Peptides were matched against the human UniProt database (20230710, 20 586 sequences), with common contaminants (344 sequences) and common tags (28 sequences) appended. Carbamidomethylation of cysteine was searched as fixed modification, whereas oxidation of methionine and acetylation at protein N-termini were defined as variable modifications. Peptides with a length between 7 and 52 amino acids were considered and results were filtered for 1% FDR at the peptide spectrum match (PSM), Peptide and Protein Group Level. Quantification was performed as specified in Biognosys BGS Factory Default settings, grouping Peptides by Stripped Sequence, and performing protein inference using IDPicker. For normalization Cross-Run Normalization in Spectronaut was activated.

Spectronaut results were exported using Pivot Reports on the Protein and Peptide level and converted to Microsoft Excel files using our in-house software MS2Go. For DIA data MS2Go utilizes the python library msReport (developed at the Max Perutz Labs Proteomics Facility) for data processing. Missing values were imputed with values obtained from a log-normal distribution with a mean of 30 in msReport and statistical significance of differentially expressed proteins was determined using limma-moderated Benjamini–Hochberg-corrected two-sided t-test [30].

### Metabolomics

Metabolites were extracted from each sample by mixing 20 μl of urine supernatants with 200 μl methanol. Samples were subsequently dried down in a vacuum centrifuge and resuspended in 0.1% formic acid. Creatinine levels were determined in a targeted LC-MS/MS experiment. Normalized to the amount of creatinine determined, another aliquot of each extracted sample was evaporated and resuspended in 130 μl ACN:H2O (80:20). Samples were then centrifuged at 4°C for 10 min at 16000 g and transferred to a glass HPLC vial. 2 μl of all samples were pooled and used as a quality control (QC) sample. Samples were randomly assigned into the autosampler, and metabolites were separated on an iHILIC®-(P) Classic HPLC column (HILICON AB, 100 x 2.1 mm; 5 µm; 200 Å, Sweden) with a flow rate of 100 µl/min delivered through an Ultimate 3000 HPLC system (Thermo Fisher Scientific, Germany). The stepwise gradient started at 90% A (ACN) and took 21 min to 60% B (25 mM ammonium bicarbonate) followed by 5 min hold at 80% B and subsequent equilibration phase at 90% A with a total run time of 35 min. Sample spectra were acquired by a high-resolution tandem mass spectrometer (Q-Exactive Focus, Thermo Fisher Scientific, Germany) in full MS mode. Metabolites were ionized via electrospray ionization in polarity switching mode after HILIC separation. Ionization potential was set to +3.5/-3.0 kV, the sheet gas flow was set to 20, and an auxiliary gas flow of 5 was used. Samples were subjected to randomized analysis, flanked by a blank and a QC sample for background correction and data normalization, respectively, occurring after every set of 8 samples. QC samples were additionally measured in data-dependent and confirmation mode to obtain MS/MS spectra for identification. The obtained data set was processed by “Compound Discoverer 3.3 SP2” (Thermo Fisher Scientific). Compounds were annotated through searching against our internal mass list database which was generated with authentic standard solutions. Additional compound annotation was conducted by searching the mzCloud database.

### Lipidomics

Lipids were extracted according to the Matyash protocol [31] using 3 ml of Urinary supernatant. Internal standard mix (PE 34:0, 830456P; PS 34:0, 840028P; LPC 17:1, 855677C; SM d35:1, 860585; purchased form Avanti Polar Lipids, USA, and PC 34:0, 37-1700-7; TG 54:0, 33-1835-9; purchased from Larodan, Sweden) was added to the samples before extraction. The organic phase of the final extraction was dried in a vacuum centrifuge and resolved in 500 µl 2-propanol:MeOH:H_2_O (70:25:10, v:v:v) before injection.

Samples were analyzed with reversed phase-UHPLC (BEH-C18, 2.1x 150 mm, 1.7 µm, Waters, Milford, USA) QTOF-MS (1290 Infinity II and 6560 IM-QTOF-MS, Agilent, Waldbronn, Germany) in positive/negative ESI QTOF-only mode. For the gradient elution, an aqueous eluent A and a 2-propanol eluent B were used, both with the following additives: Ammonium acetate (10 mM), phosphoric acid (8 μM), and formic acid (0.1 vol%). The gradient started with 60% eluent A for 0.5 min, followed by a linear decrease over 8.5 min to 20% and within 13 min to 0% A. This composition was held constant for 2.5 min and then returned to initial conditions for 5 min prior to the next injection. Eluent flow was constant 150 µl/min. In positive mode 1 µl and in negative ion mode 5 µl were injected. Column temperature was 50°C. The ESI instrument parameters were in positive mode: Gas temp 300°C, flow 10 l/min, Nebulizer 50 psi, sheath gas temp 400°C, flow 12 l/min), and in negative mode: Gas temp 300°C, flow 5 l/min, Nebulizer 30 psi, sheath gas temp 350°C, flow 12 l/min). The scan source parameters in pos and neg mode were (VCap 3500, Nozzle Voltage 500 V, Fragmentor 360, Skimmer 1 and OctopoleRFPeak 750). The data were exploratively annotated using MS-DIAL and its lipidomics database. For data integration and relative quantitation we used Lipid Data Analyzer 2.8.3_2 [32].

### TCGA survival analysis

TCGA clinical data was downloaded in R Studio with the *RTCGA* and *RTCGA.clinical* packages. The clinical data was analyzed using the *survival* and *survminer* packages and plotted using the *ggsurvplot* package in ggplot2.

### Parallel reaction monitoring mass spectrometry (PRM-MS)

#### Relative peptide amount determination

Before NanoLC-MS/MS analysis, final peptide amounts were determined by separating an aliquot of each sample on an LC-UV system equipped with a monolith column (Thermo scientific technical note 72602) and normalizing it to the peak area of 100 ng of Pierce HeLa protein digest standard (PN 88329; ThermoFisher Scientific).

#### NanoLC-MS/MS analysis

The nano HPLC system used was a Vanquish Neo UHPLC-System coupled to an Orbitrap Exploris 480 mass spectrometer, equipped with an Easy spray Source TNG (Thermo Fisher Scientific). Peptides were loaded onto a trap column (PepMap C18, 5 mm × 300 μm ID, 5 μm particles, 100 Å pore size, Thermo Fisher Scientific) by using 0.1% TFA. The trap column was switched in line with the analytical column (Double nanoViper™ PepMap C18, 500 mm × 75 μm ID, 2 μm, 100 Å, Thermo Fisher Scientific). The analytical column was connected to PepSep sprayer 1 (Bruker) equipped with a 10 μm ID fused silica electrospray emitter with an integrated liquid junction (Bruker, PN 1893527). Electrospray voltage was set to 2.3 kV. The analytical column flow was run at 230 nL/min, at 60 min binary gradient, with two mobile phases: water with 0.1% formic acid (A) and water with 80% acetonitrile and 0.08% formic acid (B). A and B were applied in linear gradients as follows (only B percentages reported): starting from 2% B: 2%-10% B in 10 min, 10%-24% B in 35 min, 24%-35% B in 15 min, 35%-95% B in 1 min, 95% for 5 min, and finally the column was equilibrated in 2% B for 3 analytical column volumes at 30°C (all % values are v/v; Water and ACN solvents were purchased from Thermo Scientific Price at LC-MS grade).

The Orbitrap Exploris 480 mass spectrometer was operated by a mixed MS method which consisted of one full scan (m/z range 380-1500; 15000 resolution; target value 100%) followed by the PRM of targeted peptides from an inclusion list (isolation window 0.8 m/z; normalized collision energy (NCE) 34; 30000 resolution, AGC target 200%). Spectra of unique peptides of the proteins of interest was recorded. Per protein at least 2 unique peptides were measured. The maximum injection time was set to 125 ms. Each precursor was measured in a 5 min time window. Peptides included in the PRM method are listed in **Supplementary Table 6.** A scheduled PRM method (sPRM) development, data processing and manual evaluation of results was performed in Skyline [33] (64-bit, v22.2.0.351). To derive the PRM score for each protein of interest (POI), the area of each identified peptide (Peptide area, PA) was summed and divided by the sum of all areas from all identified peptides from the three normalization proteins using the following formula:

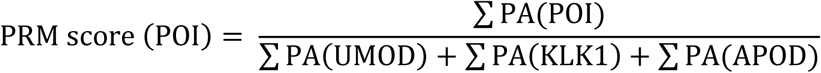

The UrineScore was calculated in the following way: The PRM score of the three POIs was calculated for controls, nccRCC, and ccRCC patients. The 95% confidence interval (CI) of the median of the PRM scores was calculated for the three POIs for the controls. (i) Haptoglobin 95% CI of median = 0.003772 – 0.03508, (ii) SAA1 95% CI of median = 0 – 5.35E-9, (iii) LCN15 95% CI of median = 0 – 2.02E-5. Then, for each control and patient sample, the PRM scores for each POI was compared to the corresponding proteins 95% CI of the controls. If the PRM score was higher than the upper limit of the controls 95% CI, a value of 1 was attributed. This calculation was done for all three proteins and the values added together, meaning that each control and patient sample will receive an UrineScore which is an integer between 0-3.

### Statistical analysis

All statistical analysis were performed in Prism 10 (GraphPad) or in RStudio (Posit). When comparing large omics datasets (proteomics, lipidomics or metabolomics) p values were calculated with limma-moderated Benjamini–Hochberg-corrected two-sided t-test after data processing for proteomics and metabolomics, and Benjamini-Hochberg adjusted. For comparisons of individual markers between groups, the distribution of the data was initially determined by Shapiro-Wilk normality test. All individual marker comparisons shown in the paper did not pass the Shapiro-Wilk normality, and subsequent analysis was either performed using Mann-Whitney U-tests (two groups) or Kruskal-Wallis test with Kruskal-Wallis test with Dunn’s multiple comparisons test (three groups). The performance of receiver operating characteristic curve (ROC) was assessed with the Area under ROC method (AUC) and a p value calculated by testing the null hypothesis that the AUC is equal to 0.5. Gene Ontology analysis was performed using online portals, Enrichr for proteomics data and MetaboAnalyst for metabolomics data.

### Author contribution

G.J. and J.M.P conceived the study. G.J. performed and designed experiments with input and help from all co-authors as follows: M.H. with mass spectrometry sample preparation and overall data analysis; T.O. with mass spectrometry; U.L, Z.K, B.E. and M.S. with ethical approval and patient sample collection; S.M. with data analysis and interpretation; M.N. with bioinformatics; G.G. and T.K. with metabolomics; G.K. and K.S. with PRM-MS; G.N.R and T.Z. with lipidomics. G.J. and J.M.P wrote the paper with input from all authors.

## Supporting information

Supplementary Table 1 Patient info

Supplementary Table 2 Supernatant proteomics

Supplementary Table 3 Sediment proteomics

Supplementary Table 4 Metabolomics

Supplementary Table 5 Lipidomics

Supplementary Table 6 PRM peptides

## Acknowledgements

Metabolomics was performed at the VBCF Metabolomics Facility which is funded by the City of Vienna through the Vienna Business Agency. Proteomics analyses were performed by the Proteomics Facility at IMP/IMBA/GMI using the VBCF instrument pool. G.J. is supported by a DOC fellowship from the Austrian Academy of Sciences. S.M. received funding from the European Union’s Horizon 2020 research and innovation programme under the Marie Sklodowska-Curie grant agreement No 841319 and the ESPRIT-Programme of the Austrian Science Fund (FWF, Project number: ESP 166). J.M.P. received funding from the Medical University of Vienna, the Austrian Academy of Sciences, the T. von Zastrow foundation, the Fundacio La Marato de TV3 (202125-31), and the Canada 150 Research Chairs Program F18-01336. We also gratefully acknowledge funding by the German Federal Ministry of Education and Research (BMBF) under the project “Microbial Stargazing - Erforschung von Resilienzmechanismen von Mikroben und Menschen” (Ref. 01KX2324).

## Data availability

All analyzed omics data (proteomics from urine supernatants, proteomics from urine sediments, lipidomics and metabolomics) are available as supplementary tables with the publication. Raw mass spectrometry proteomics data for urine supernatants, urine sediments and PRM-MS will be deposited into the PRIDE database upon favorable peer-review and made available at the date of publication.

## Supplementary tables

**Supplementary Table 1. Patient information.** Table containing information regarding patient age, gender, clinical parameters (e.g. C-reactive protein levels in serum), type of tumor and stage of tumor at the timepoint of urine collection.

**Supplementary Table 2. Supernatant proteomics.** Table containing raw expression protein data (normalized area) for each detected protein in the urine supernatant for each sample.

**Supplementary Table 3. Sediment proteomics.** Table containing raw expression protein data (normalized area) for each detected protein in the urine sediment for each sample, as well as group averages and statistics.

**Supplementary Table 4. Metabolomics.** Table containing raw expression metabolite data (normalized area) for each detected metabolite in the urine for each sample.

**Supplementary Table 5. Lipidomics.** Table containing raw expression lipid data (normalized area) for each detected lipid class and species in the urine for each sample.

**Supplementary Table 6. PRM peptides.** Table containing peptide sequences for the diagnostic proteins (SAA1, HP and LCN15) and the normalization proteins (UMOD, KLK1 and APOD) used in the PRM-MS approach.

---

**Supplementary Figure 1.**
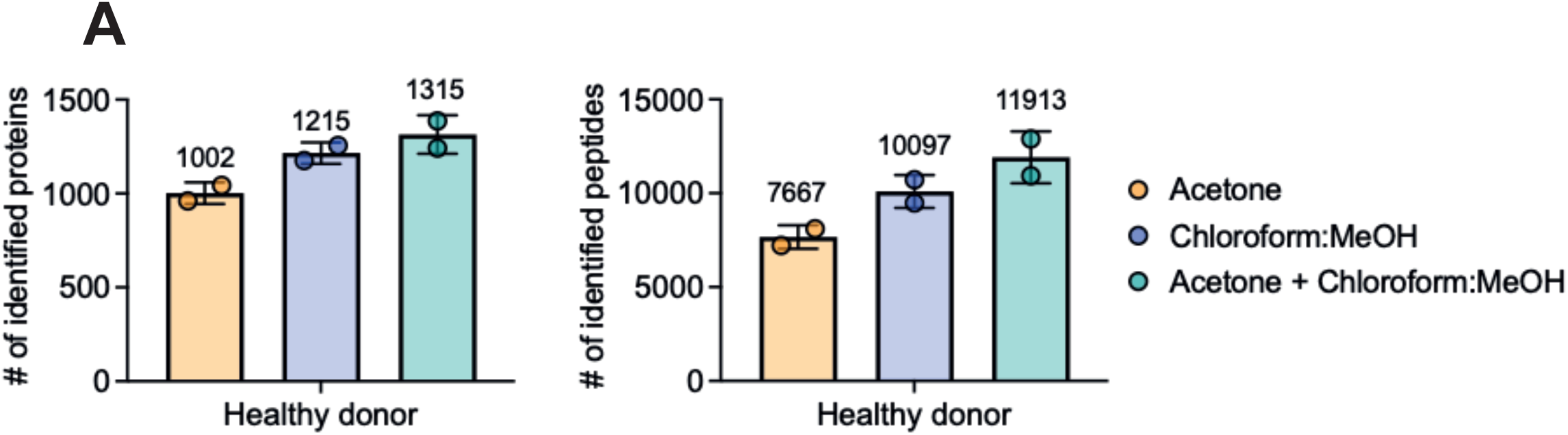
Double protein precipitation increases unique peptide and protein yields from urine supernatant. Number of identified unique proteins (left) and unique peptides (right) in a healthy control urine supernatant sample using three different protein precipitation methods. (i) Precipitation only with acetone, (ii) precipitation only with chloroform and methanol, and (iii) double protein precipitation with acetone followed by chloroform and methanol. Two technical replicates per sample is shown. MeOH = Methanol.

**Supplementary Figure 2.**
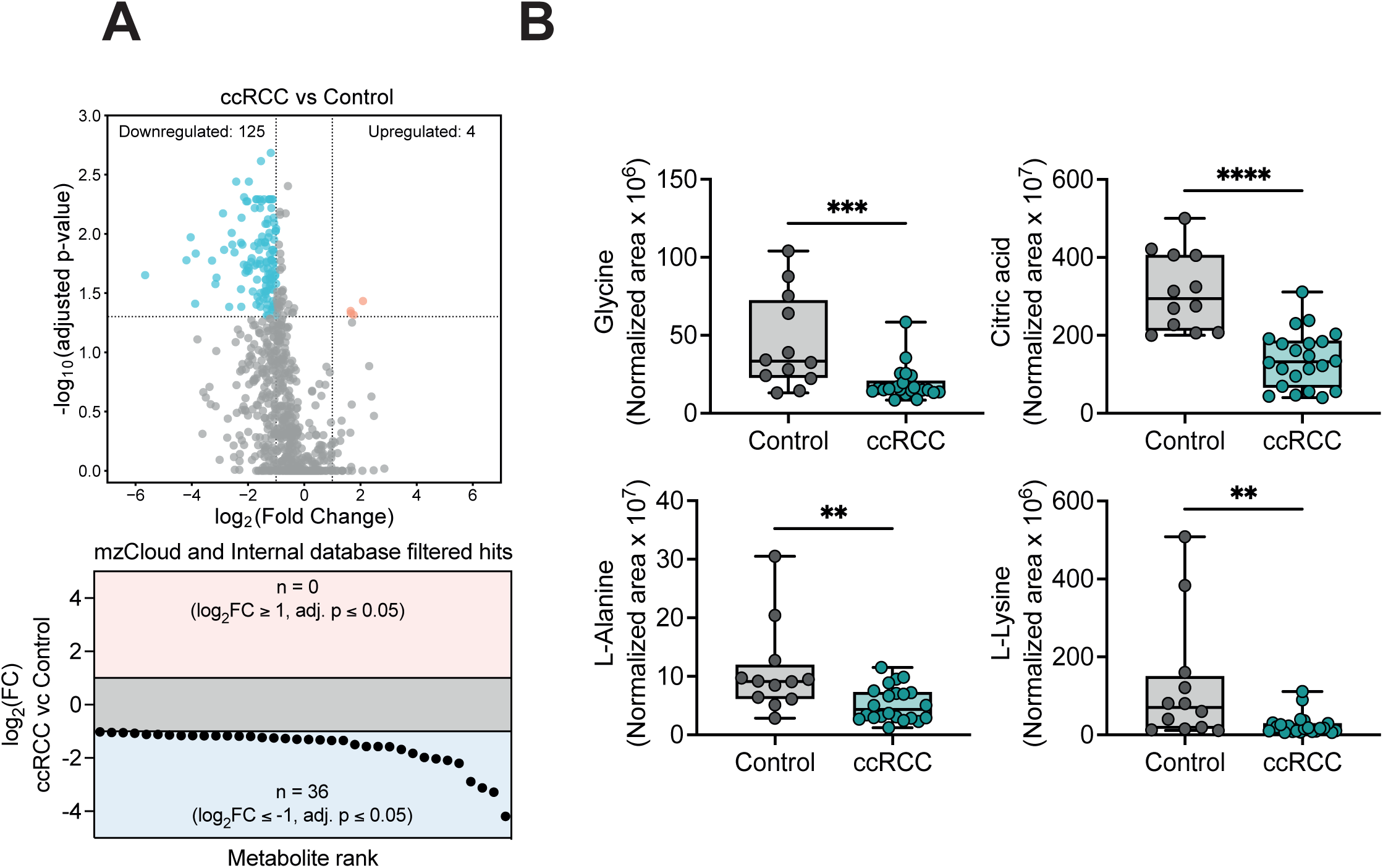
Metabolites in the urine of ccRCC patients. **A.** Volcano plot showing upregulated (red) and downregulated (blue) metabolites in ccRCC patients compared to controls. Adjusted p values calculated via limma-moderated Benjamini– Hochberg-corrected two-sided t-test. FC = fold change. **B.** Waterfall plot of significantly metabolites remaining when comparing ccRCC and control urine after filtering all hits from **A** through the mzCloud and an internal database. 36 significantly downregulated metabolites remain. **C.** Boxplots of 4 representative metabolites expression values (Glycine, Citric acid, L-Alanine and L-Lysine) as determined by metabolomics. ** = p < 0.01, *** = p < 0.001, **** = p < 0.0001.

**Supplementary Figure 3.**
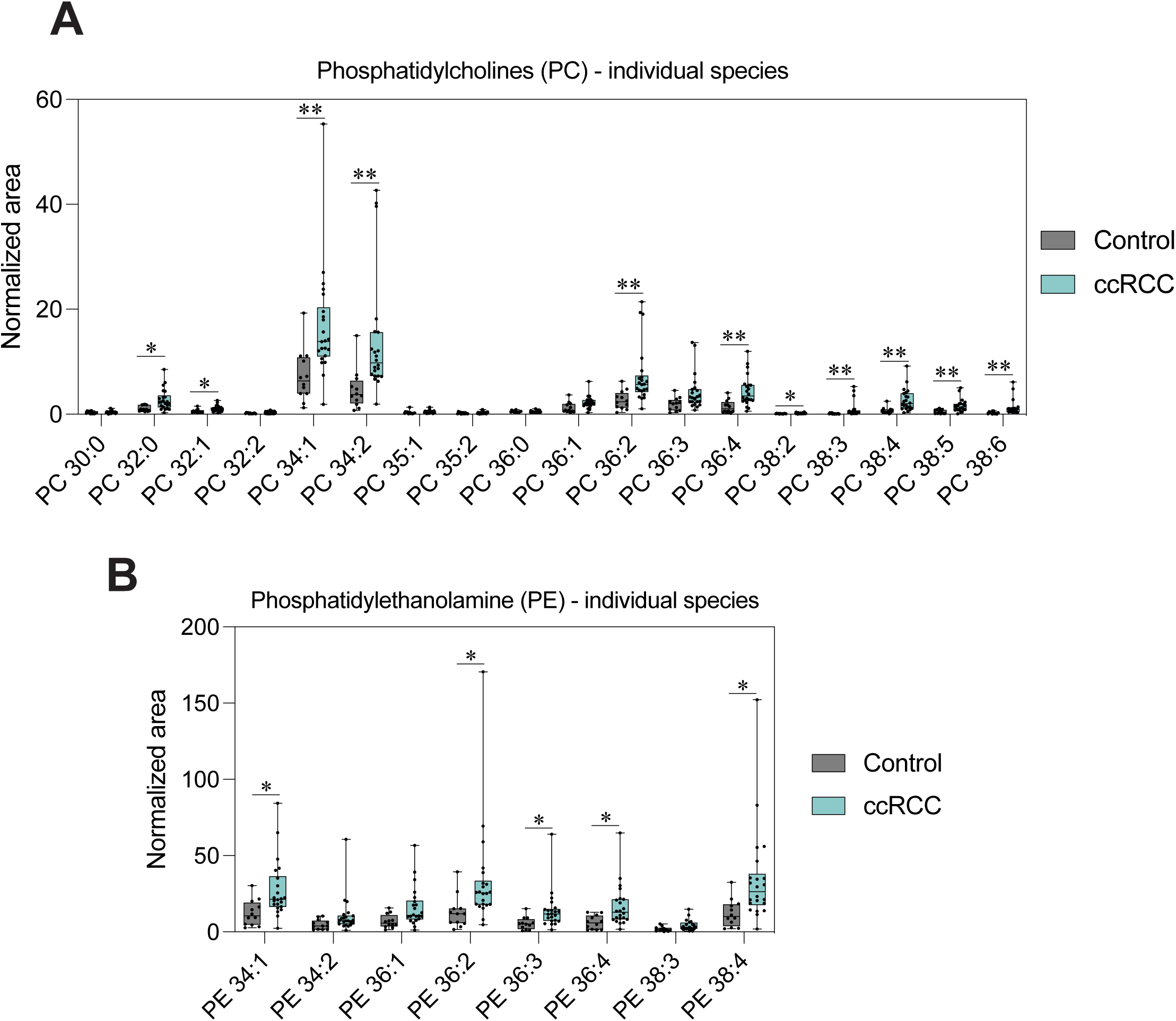
PC and PE lipid species are enriched in ccRCC urine. **A.** Boxplots of individual lipid species from the PC family. PC = Phosphatidylcholines. * = p < 0.05, ** = p < 0.01. **B.** Boxplots of individual lipid species from the PE family. PE = Phosphatidylethanolamine. * = p < 0.05.

**Supplementary Figure 4.**
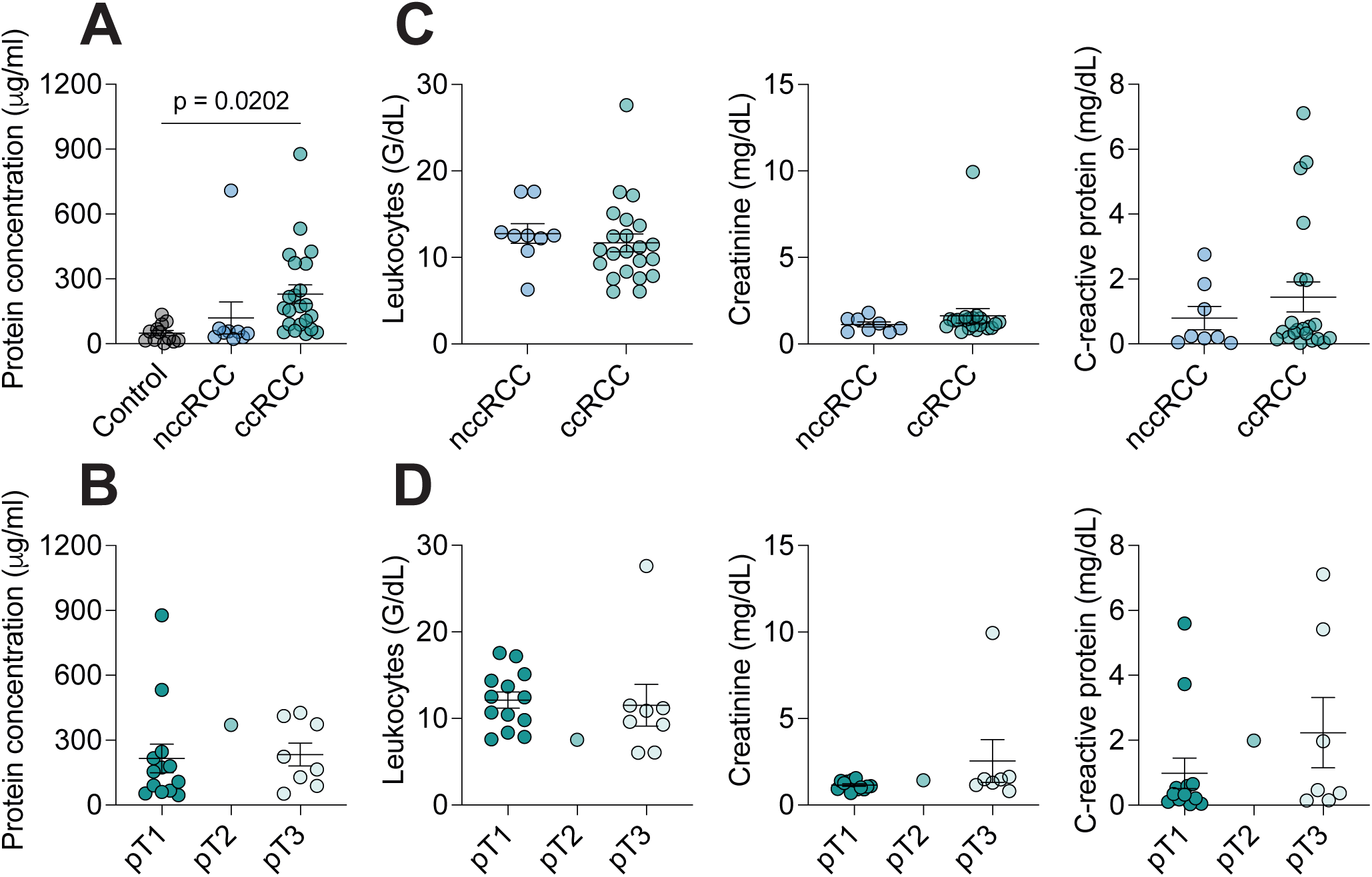
Clinical cohort overview data. **A.** Protein concentration in urine supernatant from control, nccRCC patients and ccRCC patients determined via Bradford protein assay. * = p < 0.005 **B.** Protein concentration in urine supernatant from ccRCC patients divided into pT1, pT2 and pT3 stages determined via Bradford protein assay. Abbreviations: pT1 = Tumor stage 1. The tumor is a maximum of 7 cm across. pT2 = Tumor stage 2. The tumor is larger than 7 cm across. pT3 = Tumor stage 3. The tumor has grown into a major renal vein (e.g. vena cava or renal vein) or into neighboring tissues but has not spread past Gerota’s fascia or into the adrenal gland. **C.** Leukocyte, Creatinine and C-reactive protein levels in serum, respectively, for nccRCC patients and ccRCC patients **D.** Leukocyte, Creatinine and C-reactive protein levels in serum, respectively, ccRCC patients divided into pT1, pT2 and pT3 stages. Abbreviations: pT1 = Tumor stage 1. The tumor is a maximum of 7 cm across. pT2 = Tumor stage 2. The tumor is larger than 7 cm across. pT3 = Tumor stage 3. The tumor has grown into a major renal vein (e.g. vena cava or renal vein) or into neighboring tissues but has not spread past Gerota’s fascia or into the adrenal gland.

**Supplementary Figure 5.**
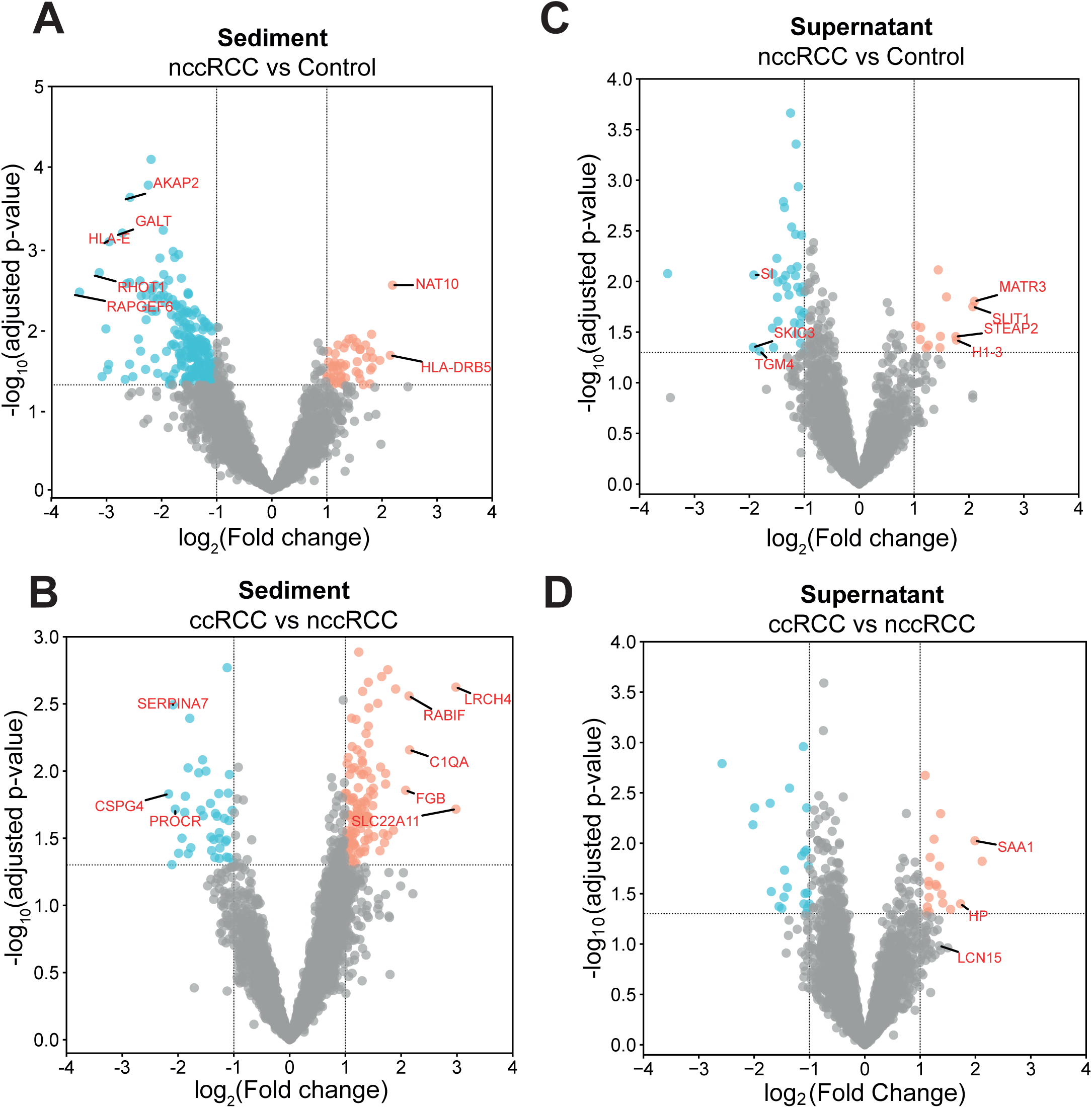
Distinct urinary protein landscapes in ccRCC and nccRCC patients. **A.** Volcano plot showing upregulated (red) and downregulated (blue) proteins in nccRCC patients compared to Controls in urine sediment. Adjusted p values calculated via limma-moderated Benjamini–Hochberg-corrected two-sided t-test. **B.** Volcano plot showing upregulated (red) and downregulated (blue) proteins in nccRCC patients compared to ccRCC patients in urine sediment. Adjusted p values calculated via limma-moderated Benjamini–Hochberg-corrected two-sided t-test. **C.** Volcano plot showing upregulated (red) and downregulated (blue) proteins in nccRCC patients compared to controls in urine supernatant. Adjusted p values calculated via limma-moderated Benjamini–Hochberg-corrected two-sided t-test. FC = fold change. **E.** Volcano plot showing upregulated (red) and downregulated (blue) proteins in ccRCC patients compared to nccRCC patients in urine supernatant. Adjusted p values calculated via limma-moderated Benjamini–Hochberg-corrected two-sided t-test. FC = fold change.

**Supplementary Figure 6.**
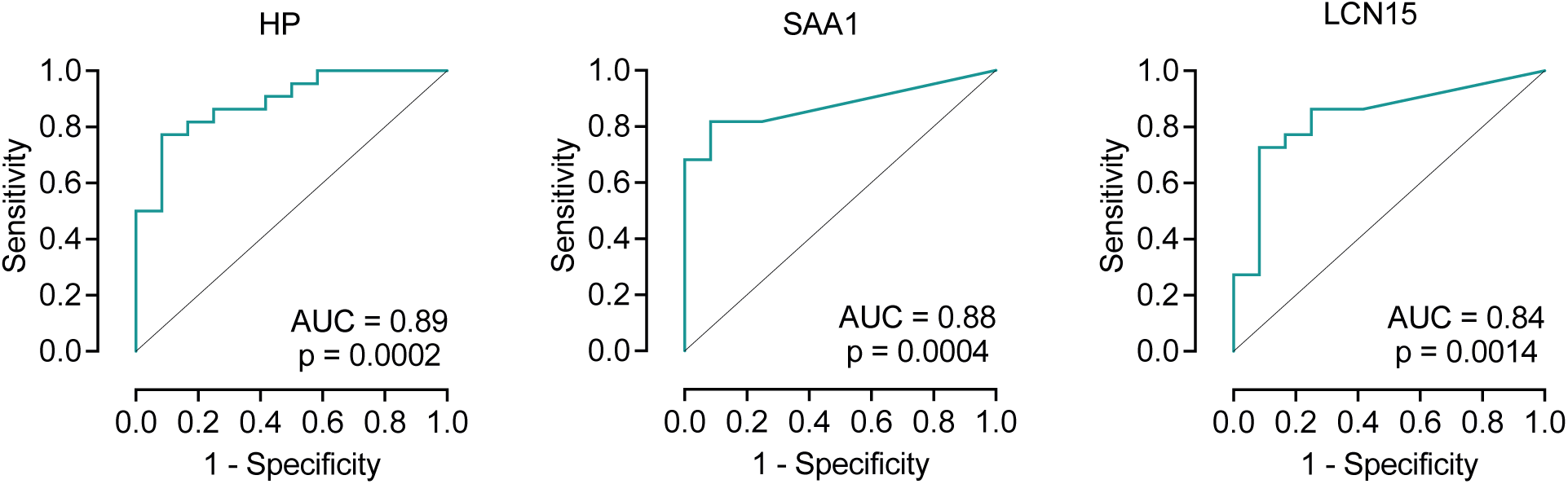
PRM-MS enables detection of SAA1, HP and LCN15 in the urine of ccRCC patients. Receiver operating characteristic curve to assess the performance of the PRM score of Haptoglobin, SAA1 and LCN15, respectively, in differentiating between control and ccRCC samples. Performance quantified through Area under the receiver operating characteristic curve (AUC) analysis.

## References

1. Hsieh, J.J., et al., Renal cell carcinoma. Nature reviews Disease primers, 2017. 3(1): p. 1–19.

2. Muglia, V.F. and A. Prando, Renal cell carcinoma: histological classification and correlation with imaging findings. Radiologia brasileira, 2015. 48: p. 166–174.

3. Giaccia, A., B.G. Siim, and R.S. Johnson, HIF-1 as a target for drug development. Nature reviews Drug discovery, 2003. 2(10): p. 803–811.

4. Lonser, R.R., et al., von Hippel-Lindau disease. The Lancet, 2003. 361(9374): p. 2059–2067.

5. Sato, Y., et al., Integrated molecular analysis of clear-cell renal cell carcinoma. Nature genetics, 2013. 45(8): p. 860–867.

6. Hakimi, A.A., et al., Clinical and pathologic impact of select chromatin-modulating tumor suppressors in clear cell renal cell carcinoma. European urology, 2013. 63(5): p. 848–854.

7. Albiges, L., et al., Pembrolizumab plus lenvatinib as first-line therapy for advanced non-clear-cell renal cell carcinoma (KEYNOTE-B61): A single-arm, multicentre, phase 2 trial. The Lancet Oncology, 2023. 24(8): p. 881–891.

8. Powles, T., et al., Pembrolizumab versus placebo as post-nephrectomy adjuvant therapy for clear cell renal cell carcinoma (KEYNOTE-564): 30-month follow-up analysis of a multicentre, randomised, double-blind, placebo-controlled, phase 3 trial. The Lancet Oncology, 2022. 23(9): p. 1133–1144.

9. Vano, Y.-A., et al., Nivolumab, nivolumab–ipilimumab, and VEGFR-tyrosine kinase inhibitors as first-line treatment for metastatic clear-cell renal cell carcinoma (BIONIKK): A biomarker-driven, open-label, non-comparative, randomised, phase 2 trial. The Lancet Oncology, 2022. 23(5): p. 612–624.

10. Gray, R.E. and G.T. Harris, Renal cell carcinoma: diagnosis and management. American family physician, 2019. 99(3): p. 179–184.

11. Burg, M., et al., Organic solutes in fluid absorption by renal proximal convoluted tubules. American Journal of Physiology-Legacy Content, 1976. 231(2): p. 627–637.

12. Lucarelli, G., et al., Metabolomic insights into pathophysiological mechanisms and biomarker discovery in clear cell renal cell carcinoma. Expert review of molecular diagnostics, 2019. 19(5): p. 397–407.

13. Zhang, Y., et al., Single-cell analyses of renal cell cancers reveal insights into tumor microenvironment, cell of origin, and therapy response. Proceedings of the National Academy of Sciences, 2021. 118(24): p. e2103240118.

14. Li, M., et al., Liquid biopsy at the frontier in renal cell carcinoma: recent analysis of techniques and clinical application. Molecular Cancer, 2023. 22(1): p. 37.

15. Farber, N.J., et al., Renal cell carcinoma: the search for a reliable biomarker. Translational cancer research, 2017. 6(3): p. 620.

16. Rustam, Y.H. and G.E. Reid, Analytical challenges and recent advances in mass spectrometry based lipidomics. Analytical chemistry, 2018. 90(1): p. 374–397.

17. Peterson, A.C., et al., Parallel reaction monitoring for high resolution and high mass accuracy quantitative, targeted proteomics. Molecular & cellular proteomics, 2012. 11(11): p. 1475–1488.

18. Bukavina, L., et al., Epidemiology of Renal Cell Carcinoma: 2022 Update. Eur Urol, 2022. 82(5): p. 529–542.

19. Diana, P., et al., Screening programs for renal cell carcinoma: a systematic review by the EAU young academic urologists renal cancer working group. World journal of urology, 2023. 41(4): p. 929–940.

20. Usher-Smith, J.A., et al., The Yorkshire Kidney Screening Trial (YKST): protocol for a feasibility study of adding non-contrast abdominal CT scanning to screen for kidney cancer and other abdominal pathology within a trial of community-based CT screening for lung cancer. BMJ open, 2022. 12(9): p. e063018.

21. Du, W., et al., HIF drives lipid deposition and cancer in ccRCC via repression of fatty acid metabolism. Nature communications, 2017. 8(1): p. 1769.

22. Marques, L.R., et al., Reverse cholesterol transport: molecular mechanisms and the non-medical approach to enhance HDL cholesterol. Frontiers in Physiology, 2018. 9: p. 526.

23. Sandim, V., et al. Proteomic analysis reveals differentially secreted proteins in the urine from patients with clear cell renal cell carcinoma. in Urologic Oncology: Seminars and Original Investigations. 2016. Elsevier.

24. Santorelli, L., et al., In-depth mapping of the urinary N-glycoproteome: distinct signatures of ccRCC-related progression. Cancers, 2020. 12(1): p. 239.

25. Chinello, C., et al., Proteomics of liquid biopsies: Depicting RCC infiltration into the renal vein by MS analysis of urine and plasma. Journal of proteomics, 2019. 191: p. 29–37.

26. Yang, Y., et al., Excavation of diagnostic biomarkers and construction of prognostic model for clear cell renal cell carcinoma based on urine proteomics. Frontiers in Oncology, 2023. 13: p. 1170567.

27. Di Meo, A., et al., Searching for prognostic biomarkers for small renal masses in the urinary proteome. International journal of cancer, 2020. 146(8): p. 2315–2325.

28. Wessel, D. and U. Flügge, A method for the quantitative recovery of protein in dilute solution in the presence of detergents and lipids. Analytical biochemistry, 1984. 138(1): p. 141–143.

29. Bruderer, R., et al., Extending the limits of quantitative proteome profiling with data-independent acquisition and application to acetaminophen-treated three-dimensional liver microtissues*[S]. Molecular & Cellular Proteomics, 2015. 14(5): p. 1400–1410.

30. Smyth, G.K., Linear models and empirical bayes methods for assessing differential expression in microarray experiments. Statistical applications in genetics and molecular biology, 2004. 3(1).

31. Matyash, V., et al., Lipid extraction by methyl-tert-butyl ether for high-throughput lipidomics. Journal of lipid research, 2008. 49(5): p. 1137–1146.

32. Hartler, J., et al., Deciphering lipid structures based on platform-independent decision rules. Nature methods, 2017. 14(12): p. 1171–1174.

33. MacLean, B., et al., Skyline: an open source document editor for creating and analyzing targeted proteomics experiments. Bioinformatics, 2010. 26(7): p. 966–968.

